# Beyond Marginal Coverage: Class-Conditional Conformal Prediction Reveals Diagnosis-Specific Uncertainty Structure in Psychiatric Neuroimaging

**DOI:** 10.64898/2026.09.23.753958

**Authors:** Hooman Rokham, Haleh Falakshahi, Vince D. Calhoun

## Abstract

Conformal prediction is increasingly proposed for clinical decision support because it provides distribution-free coverage guarantees that hold for any model and any data distribution. The guarantee routinely reported, however, is *marginal*, and we show that in multi-class diagnostic classification it can be satisfied exactly while individual diagnoses are covered very unevenly. On a four-way mood and psychosis classification task over 1,520 subjects from three studies and 14 acquisition sites, marginal split-conformal calibration attained 0.9000 empirical coverage against a nominal 0.90, while healthy controls were covered at 0.941 and schizoaffective disorder at 0.819. The surplus and the deficit cancel, so the aggregate figure is uninformative about either. Class-conditional (Mondrian) calibration reduced this 12.2-point disparity to 0.4 points at a cost of 0.07 labels in mean set size, under 3%.

Holding coverage fixed across diagnoses has a further consequence: the residual variation in set size can no longer be attributed to class prior or per-class accuracy, and becomes interpretable as a property of the subject. We show that the resulting prediction sets separate subjects into confident, boundary, ambiguous and unresolved strata, and that the proportion of subjects independently flagged as label-ambiguous by a structural-MRI model trained separately on the same cohort rises monotonically across these strata (34.7%, 57.0%, 68.0%, 82.1%; *p* = 2.1 × 10^*−*17^). No schizoaffective subject and 0.9% of bipolar subjects reach the confident stratum, against 18.4% of controls and 13.0% of schizophrenia subjects.

Set size at matched coverage also provides a comparison between representations that accuracy cannot. Applied to structural MRI, functional MRI and their fusion, it reveals a sign reversal that accuracy conceals: fusion reduces set size for bipolar (−0.25) and schizoaffective (−0.23) subjects and increases it for controls (+0.16), and the aggregate fusion benefit itself changes sign below *α* = 0.10, while top-1 accuracy rises monotonically from 0.495 to 0.578 to 0.611 across the three models. Reporting marginal coverage alone is insufficient whenever diagnostic classes are unbalanced.

## 1 Introduction

Deep classifiers are increasingly applied to biomedical signals, yet they are rarely deployed with calibrated statements of uncertainty. Conformal prediction offers an attractive remedy: given any pre-trained model and a held-out calibration set, it returns a *set* of labels guaranteed to contain the truth with probability at least 1− *α*, in finite samples and without assumptions on the data distribution. The guarantee requires only that calibration and test scores be exchangeable, which makes it unusually easy to adopt, and unusually easy to report without checking.

The guarantee that split conformal prediction actually delivers is *marginal*: the probability is taken over the joint draw of covariates and label. It therefore constrains only an average over the label distribution and permits coverage to be distributed unevenly across classes. A single threshold will over-cover classes that are abundant and well separated and under-cover classes that are rare or confusable, and these two errors can cancel exactly in the aggregate. Where classes are balanced and well separated, marginal and class-conditional coverage nearly coincide and the choice between them has little consequence. Neither condition holds in psychiatric neuroimaging: diagnostic groups differ in size by a factor of two or more, and the boundaries between them are contested on clinical grounds. Schizoaffective disorder in particular is defined as an intermediate presentation between schizophrenia and the mood disorders, and its status as a distinct category has been debated since its introduction.

Split conformal prediction was formalized by Vovk et al. [1] and analyzed by Lei et al. [2]. Common score functions include the least ambiguous set-valued classifier (LAC) [3], adaptive prediction sets (APS) [4] and regularized APS (RAPS) [5]. Class-conditional, or Mondrian, calibration [6, 7] restores per-class validity, and related work addresses covariate shift [8] and label noise [9]. Applications in neuroimaging remain scarce, and where conformal methods are used, coverage is typically asserted from the nominal *α* rather than measured on held-out data. We therefore measure coverage directly, on a four-way mood and psychosis classification task in which unbalanced groups and contested boundaries are both present.

Our contributions are fourfold. First, we quantify the disparity empirically on 1,520 subjects: marginal calibration attains its nominal level to within 10^−4^ while per-class coverage differs by 12.2 percentage points. Second, we show that class-conditional calibration removes the disparity at an efficiency cost below 3%. Third, we show that the resulting prediction sets induce an interpretable stratification of subjects into confident, boundary, ambiguous and unresolved cases, validated against an independently trained model on a different imaging modality. Fourth, we use set size at matched coverage to compare structural MRI, functional MRI and their fusion, and find that fusion is informative at the mood–psychosis boundary and detrimental for controls, a reversal that accuracy records as a uniform improvement.

## 2 Methods

### 2.1 Split conformal classification

Let 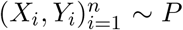 with *Y*_*i*_ ∈ Y = {1, …, *K*} and let 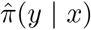 be the softmax output of a classifier trained on a disjoint set. For a nonconformity score *s*(*x, y*), split conformal prediction computes

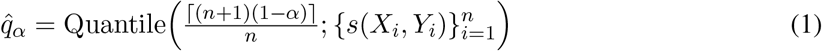

on a calibration set and returns 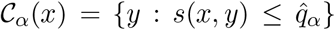. If the calibration and test scores are exchangeable,

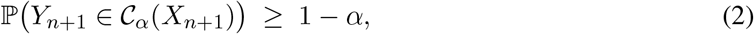

with no assumption on *P* or on the classifier. The (*n* + 1) correction in (1) is what makes the bound finite-sample exact rather than asymptotic.

### 2.2 Marginal versus class-conditional validity

The probability in (2) is taken over the joint draw of (*X*_*n*+1_, *Y*_*n*+1_), so it constrains only the average over the label distribution. A single threshold may therefore allocate excess coverage to classes that are abundant and well separated, financed by a deficit on classes that are rare or confusable, while the reported marginal number remains exactly at its nominal level. Section 4 demonstrates that the two effects cancel to within 10^−4^ in our data while individual classes differ by more than twelve points.

A second consequence is interpretive. Under a single threshold, |C_*α*_(*x*)| is monotone in the rank the classifier assigns to the true label, so mean set size within a class is largely determined by that class’s accuracy and prior. Any claim that a diagnostic category is “intrinsically more uncertain” therefore requires coverage to be equalized first, and external evidence thereafter. This ordering of operations is what makes Sections 4.3 and 4.4 interpretable rather than circular.

### 2.3 Class-conditional calibration

We compute a separate threshold per class [6, 7],

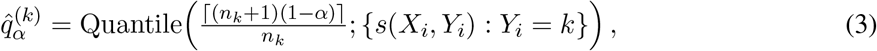

and form 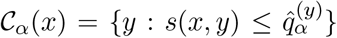, giving ℙ (*Y* ∈ C_*α*_(*X*) | *Y* = *k*) ≥ 1 − *α* for every *k*. This requires *n*_*k*_ ≥ ⌈1*/α*⌉ − 1 calibration points per class; our smallest group (*n* = 234, 117 after the split) admits *α* ≥ 0.009.

### 2.4 Nonconformity scores

We compare three scores. LAC [3] takes 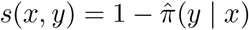, which yields the smallest average sets but the weakest conditional coverage. APS [4] accumulates sorted probability mass,

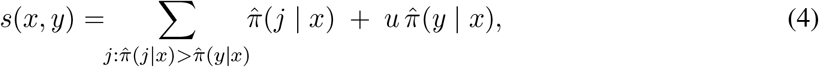

with *u* ~ Unif(0, 1) drawn once per sample; the randomization is what makes coverage exact rather than conservative. RAPS [5] adds *λ* max(0, rank(*y*) − *k*_reg_) to (4) to suppress the deep tail, with *λ* = 0.05 and *k*_reg_ = 1 here. We report all three and use APS for the analyses that follow, for comparability with prior work.

### 2.5 Evaluation

Beyond marginal coverage we report per-class coverage and mean set size. We also report residual ambiguity, E[log_2_ |C_*α*_(*X*)|] in bits, which is comparable across representations at matched coverage in a way that accuracy is not, and size-stratified coverage, which exposes the case where marginal coverage is met while the hardest stratum is badly under-covered. All calibration and evaluation splits are drawn at the *subject* level and repeated 300 to 400 times; we report the mean and standard deviation across splits.

## 3 Data and Experimental Setup

We use resting-state fMRI from 1,520 subjects pooled across three studies, FBIRN [13] and the first two waves of B-SNIP [12], spanning 14 acquisition sites (Table 1). Diagnostic groups are healthy controls (NC, *n* = 566), bipolar disorder (BP, 234), schizoaffective disorder (SAD, 249) and schizophrenia (SZ, 471). We note explicitly that FBIRN contributes only NC and SZ subjects, so study membership is partially informative of the label for 286 subjects. This is a property of the pooled cohort rather than of the calibration procedure, and it affects the classifier rather than the coverage comparison, which is made within a fixed set of predictions.

**Table 1.** Cohort composition. FBIRN contributes no BP or SAD subjects.

| Study | NC | BP | SAD | SZ | Total |
| --- | --- | --- | --- | --- | --- |
| FBIRN | 145 | – | – | 141 | 286 |
| B-SNIP-1 | 216 | 145 | 117 | 173 | 651 |
| B-SNIP-2 | 205 | 89 | 132 | 157 | 583 |
| Total | 566 | 234 | 249 | 471 | 1520 |

Preprocessing used SPM12: the first five volumes were discarded, followed by slice-timing and motion correction, normalization to MNI space at 3 × 3 × 3 mm^3^, and smoothing at 6 mm FWHM. Features are the 53 intrinsic connectivity network time courses produced by the NeuroMark 1.0 ICA pipeline [11]. The structural pathway used gray-matter maps from the same subjects.

Classifiers are ensembles of convolutional networks trained with repeated five-fold cross-validation over three class-balanced subsampling sets: a fully convolutional 1D network on the ICA time courses for the functional pathway, and a 3D network on gray-matter maps for the structural pathway. Each subject receives a probability vector averaged over the five to fifteen folds in which that subject was held out, so the probabilities we calibrate are out-of-fold throughout. Four-way top-1 accuracy is 0.578 for the functional model and 0.495 for the structural model, against a majority-class baseline of 0.372.

The conformal layer is applied post hoc to these probabilities; the classifiers are neither retrained nor modified. Calibration and evaluation sets are disjoint at the *subject* level, and no subject contributes to both.

## 4 Results

### 4.1 The marginal guarantee holds; the class-wise one does not

Marginal calibration is valid exactly as the theory promises: empirical coverage is 0.9000 against a nominal 0.90 (*α* = 0.10) and 0.9494 against 0.95 (*α* = 0.05), averaged over 300 subject-level splits. Reported alone, these numbers suggest a well-calibrated system.

They are not informative about any individual diagnosis. At *α* = 0.10 (Table 2, Fig. 2a) healthy controls are covered at 0.941 and schizoaffective disorder at 0.819, a spread of 12.2 percentage points, with SAD missing its nominal level by 8.1. Expressed as error rates, the schizoaffective miscoverage rate is 0.181 against a promised 0.100, 1.81 times higher than advertised, which corresponds to approximately 20 schizoaffective subjects in this cohort missed beyond what the stated guarantee allows. Healthy controls receive approximately 23 fewer misses than promised. The surplus and the deficit sum to zero, which is precisely why the marginal figure lands on target.

**Table 2.** Coverage and efficiency at *α* = 0.10, mean over 300 subject-level splits. Marginal calibration attains its nominal level while under-covering SAD by 8.1 points.

| Class | Coverage | | Mean $\vert \mathcal{C}\vert $ | |
| --- | --- | --- | --- | --- |
|  | Marginal | Mondrian | Marginal | Mondrian |
| BP | 0.907 | 0.903 | 3.06 | 3.07 |
| NC | 0.941 | 0.901 | 2.23 | 2.35 |
| SAD | **0.819** | 0.905 | 3.09 | 3.10 |
| SZ | 0.890 | 0.902 | 2.51 | 2.57 |
| Overall | 0.900 | 0.902 | 2.58 | 2.65 |
| Spread | **0.1215** | **0.0044** |  |  |

The pattern is not confined to one operating point: Fig. 1a shows the SAD curve lying below the diagonal across the entire range *α* ∈ [0.02, 0.30], while the marginal curve tracks it almost perfectly. The ordering of per-class coverage follows class abundance and per-class accuracy, which is what a single shared threshold must produce.

**Figure 1.**
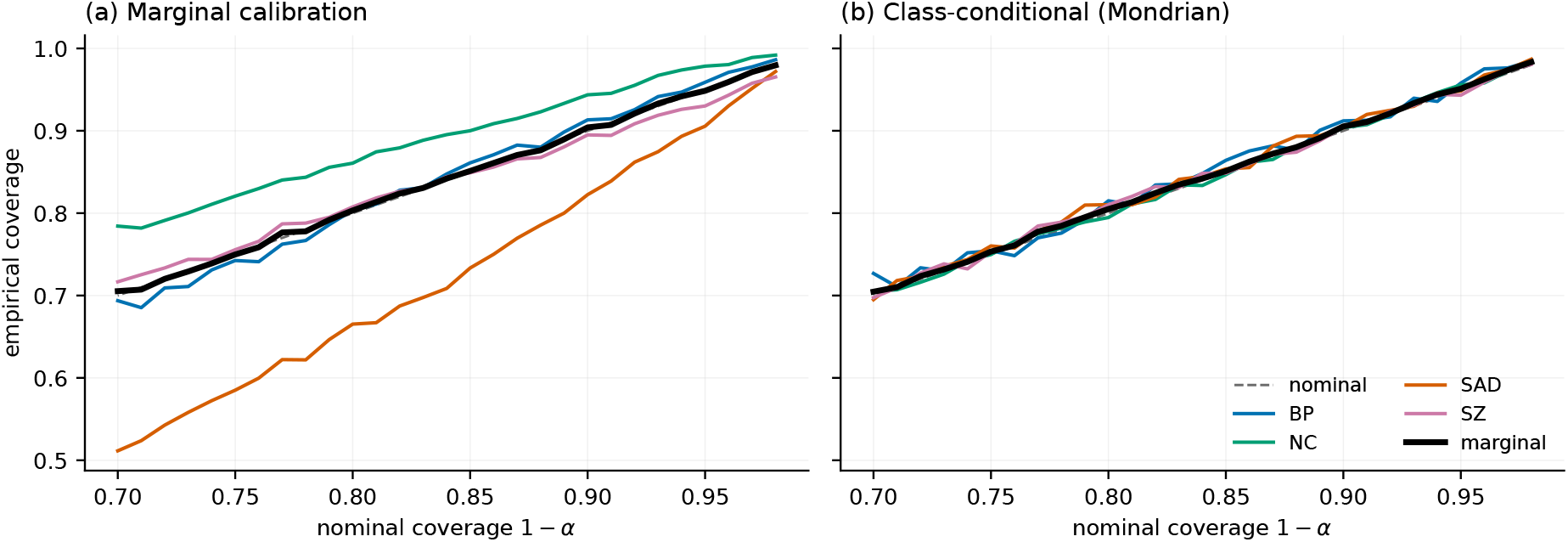
Empirical versus nominal coverage. (a) Marginal calibration: the aggregate curve (black) tracks the diagonal while SAD lies far below it throughout. (b) Class-conditional calibration collapses all four onto the diagonal.

**Figure 2.**
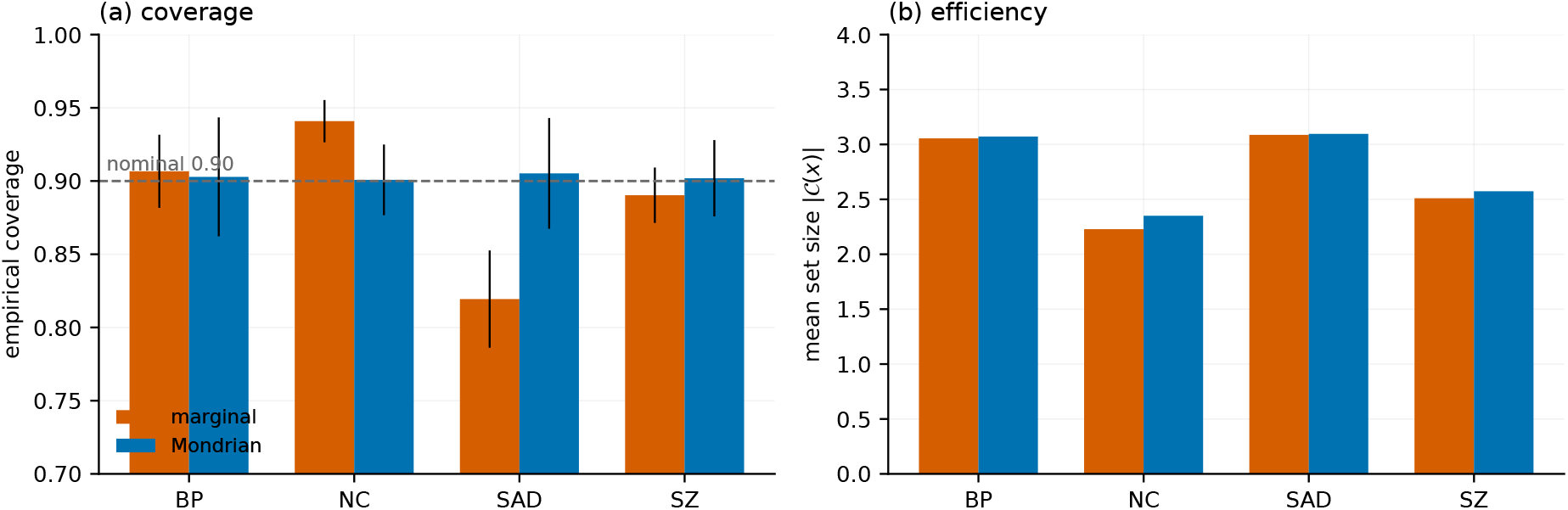
Per-class coverage and efficiency at *α* = 0.10; error bars are standard deviations over 300 splits. The coverage repair costs 0.07 labels of set size.

### 4.2 Class-conditional calibration restores per-class validity

Mondrian calibration (3) brings every class to within 0.4 percentage points of nominal (BP 0.903, NC 0.901, SAD 0.905, SZ 0.902), reducing the spread from 0.1215 to 0.0044, a 28-fold reduction. Mean set size rises from 2.582 to 2.652, an efficiency cost of 0.070 labels, or 2.7%. Across 1,520 subjects this corresponds to approximately 106 additional labels in total, or roughly seven subjects in a hundred seeing one additional diagnosis in their prediction set. Fig. 1b shows all four curves collapsed onto the diagonal.

Among score functions at *α* = 0.10 under marginal calibration, LAC is the most efficient (mean |C| = 2.48, 18.9% singletons), APS intermediate (2.59, 13.2%), and RAPS with *λ* = 0.05 the least (2.68, 0.7%); all three attain nominal marginal coverage, and all three exhibit the same per-class disparity, under-covering BP and SAD by comparable margins. The disparity is therefore a property of marginal calibration rather than of any particular score.

### 4.3 Prediction sets stratify subjects, and the strata are externally validated

Because the class-conditional procedure equalizes coverage, the remaining variation in set structure is no longer attributable to class prior or per-class accuracy. We therefore assign each subject the modal set size observed across the 400 evaluation splits and obtain four strata: *confident* (|C| ≤ 1), *boundary* (|C| = 2), *ambiguous* (|C| ≥ 3), and *unresolved*, defined as subjects whose true label is excluded from the set in a majority of splits.

To test whether these strata reflect properties of the subjects rather than of the functional classifier, we compare them against an ambiguity marker obtained from the structural-MRI model, whose per-subject cross-validated prediction consistency labels each subject reliable or ambiguous [10]. The two markers agree on 62.0% of subjects against 49.8% expected by chance given the marginals (Cohen’s *κ* = 0.24), so they are related but far from redundant, and the structural model shares no predictions with the functional conformal procedure.

The proportion of structurally flagged subjects increases monotonically across the four strata, from 34.7% to 57.0%, 68.0% and 82.1% (Table 3, Fig. 3a; *χ*^2^ test of independence, *p* = 2.1 × 10^−17^). Set size itself differs by structural marker (*p* = 8.3 × 10^−16^, Kruskal–Wallis), with mean |C| of 2.79 against 2.42 (*d* = 0.47, Fig. 3c). Two models trained on distinct tissue properties therefore agree on which subjects are difficult, which would not follow if set size merely restated the functional model’s error rate. For contrast, repeating the test with the functional model’s own marker inflates the effect to *d* = 1.03; that comparison is circular by construction, since both quantities are functions of the same fold predictions, and we report it only to bound how much of the apparent uncertainty–noise coupling in the literature may be an artifact of shared predictions.

**Table 3.** Subject strata induced by class-conditional prediction sets at *α* = 0.10. Upper panel: the proportion flagged by an independent structural-MRI model increases monotonically across strata (*p* = 2.1 × 10^*−*17^). Lower panel: distribution of each diagnosis across strata (row percentages).

| Stratum | $n$ | % of cohort | Coverage | Struct. flagged |
| --- | --- | --- | --- | --- |
| Confident | 167 | 11.0 | 0.903 | 34.7% |
| Boundary | 365 | 24.0 | 0.848 | 57.0% |
| Ambiguous | 932 | 61.3 | 0.969 | 68.0% |
| Unresolved | 56 | 3.7 | 0.161 | 82.1% |

| Diagnosis | Confident | Boundary | Ambiguous | Unresolved |
| --- | --- | --- | --- | --- |
| BP | 0.9% | 12.4% | 82.1% | 4.7% |
| NC | 18.4% | 33.0% | 45.8% | 2.8% |
| SAD | 0.0% | 10.4% | 85.5% | 4.0% |
| SZ | 13.0% | 26.1% | 56.9% | 4.0% |

**Figure 3.**
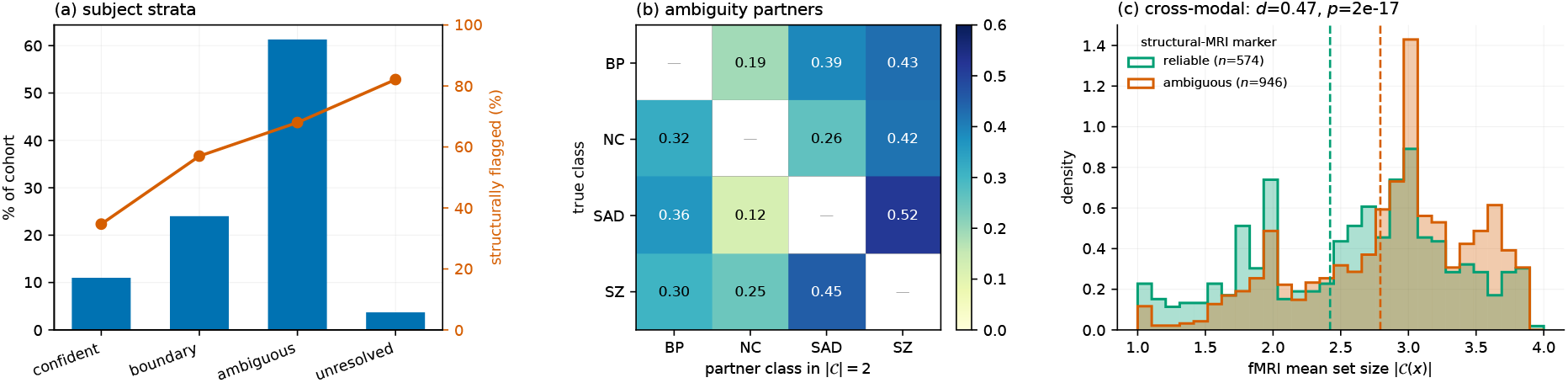
(a) Subject strata: bars give the share of the cohort, the line gives the proportion independently flagged by the structural-MRI model. (b) Given a size-two set, which class accompanies the true one (row-normalized). (c) Functional-MRI prediction-set size for subjects labelled reliable or ambiguous by the independently trained structural-MRI model.

The strata are not distributed uniformly across diagnoses (Table 3, lower panel). No schizoaffective subject and 0.9% of bipolar subjects reach the confident stratum, against 18.4% of controls and 13.0% of schizophrenia subjects; conversely 85.5% of schizoaffective and 82.1% of bipolar subjects fall in the ambiguous stratum. Confident predictions in this cohort are thus almost exclusively control or schizophrenia cases, and the two intermediate diagnoses admit essentially no confidently separable subjects.

Within the boundary stratum, the class accompanying the true label is itself informative (Fig. 3b). Schizoaffective disorder is accompanied by schizophrenia in 52% of cases and by bipolar disorder in 36%, and by healthy controls in only 12%, the smallest entry in the matrix and the smallest at every *α* we examined (0.046 at *α* = 0.05 rising to 0.186 at *α* = 0.30). Schizoaffective disorder is therefore the category least separable from the two psychosis poles and most separable from health, which is consistent with its position in the diagnostic hierarchy. The finer orderings among the remaining pairs were not stable across operating points and we do not interpret them.

The unresolved stratum comprises 56 subjects (3.7%) whose true label is recovered in only 16.1% of splits, and of whom 82.1% are also flagged structurally. These are candidates for review rather than prediction: the marginal guarantee is satisfied in aggregate while these subjects are missed systematically, and identifying them is not possible from a point prediction or from a confusion matrix.

### 4.4 Which modality is informative, and for whom

Mean set size at matched coverage provides a comparison between feature sets that accuracy cannot: because the class-conditional procedure fixes coverage at 1 − *α* for every diagnosis and every model, the remaining set size measures how much ambiguity each representation leaves in the label system. We therefore calibrated three models identically: the functional model, the structural model, and their late fusion by equal-weight averaging of softmax outputs.

At matched coverage (0.903), the structural model leaves 2.98 labels, the functional model 2.65, and the fusion 2.59 (Table 4); the functional advantage over structural is large and consistent across subjects (Δ = 0.32, *p* = 2 × 10^−48^, Wilcoxon signed-rank), while the gain from fusion over the functional model alone is small in aggregate (Δ = 0.07, *p* = 5 × 10^−11^). Expressed in bits of residual ambiguity, the three models leave 1.51, 1.31 and 1.30 bits respectively.

**Table 4.** Modality comparison at matched class-conditional coverage (*α* = 0.10). Smaller |C| indicates a representation that leaves less ambiguity in the diagnostic label system.

| Model | Acc. | Cov. | Mean $\vert \mathcal{C}\vert $ | | | |
| --- | --- | --- | --- | --- | --- | --- |
|  |  |  | All | BP | SAD | NC |
| Structural | 0.495 | 0.904 | 2.98 | 3.36 | 3.33 | 2.87 |
| Functional | 0.578 | 0.903 | 2.65 | 3.07 | 3.10 | **2.35** |
| Fusion | 0.611 | 0.903 | **2.59** | **2.81** | **2.86** | 2.51 |

The aggregate figure, however, conceals a sign reversal. Fusion reduces set size for bipolar (−0.25) and schizoaffective (−0.23) subjects and increases it for controls (+0.16); fusion shrinks the set for 73.1% of bipolar and 71.9% of schizoaffective subjects but for only 43.1% of controls, hurting 51.2% of them. Structural information is therefore informative precisely at the mood–psychosis boundary, where the functional model is least decisive, and is a liability on the class the functional model already separates. Top-1 accuracy rises monotonically from 0.495 to 0.578 to 0.611 across the three models and records none of this.

The reversal is not an artifact of the operating point (Table 5, Fig. 4). Sweeping *α* ∈ [0.02, 0.30], the fusion gain stays positive for bipolar (+0.04 to +0.26) and schizoaffective (+0.03 to +0.25) subjects and negative for controls (−0.22 to −0.02) throughout, with no crossover; only schizophrenia changes sign, below *α* = 0.05. The aggregate gain does cross zero between *α* = 0.05 and 0.10, so at strict confidence levels fusion is net harmful (−0.08 labels at *α* = 0.02), the penalty on the largest group outweighing the gain on the two smallest. Whether a second modality is worth acquiring therefore depends on the confidence level demanded, a dependence accuracy cannot express.

**Table 5.** Fusion gain, defined as mean |C| under the functional model minus mean |C| under fusion, as a function of *α*. Positive values indicate that fusion reduces ambiguity. The sign is stable for BP, SAD and NC across the whole range; the aggregate crosses zero between *α* = 0.05 and 0.10.

| $\alpha$ | BP | NC | SAD | SZ | All |
| --- | --- | --- | --- | --- | --- |
| 0.02 | +0.044 | −0.194 | +0.026 | −0.051 | −0.077 |
| 0.05 | +0.138 | −0.217 | +0.135 | +0.068 | −0.016 |
| 0.10 | +0.251 | −0.158 | +0.225 | +0.153 | +0.064 |
| 0.15 | +0.256 | −0.125 | +0.250 | +0.173 | +0.087 |
| 0.20 | +0.246 | −0.091 | +0.247 | +0.195 | +0.105 |
| 0.25 | +0.229 | −0.052 | +0.245 | +0.191 | +0.115 |
| 0.30 | +0.228 | −0.016 | +0.246 | +0.186 | +0.127 |

**Figure 4.**
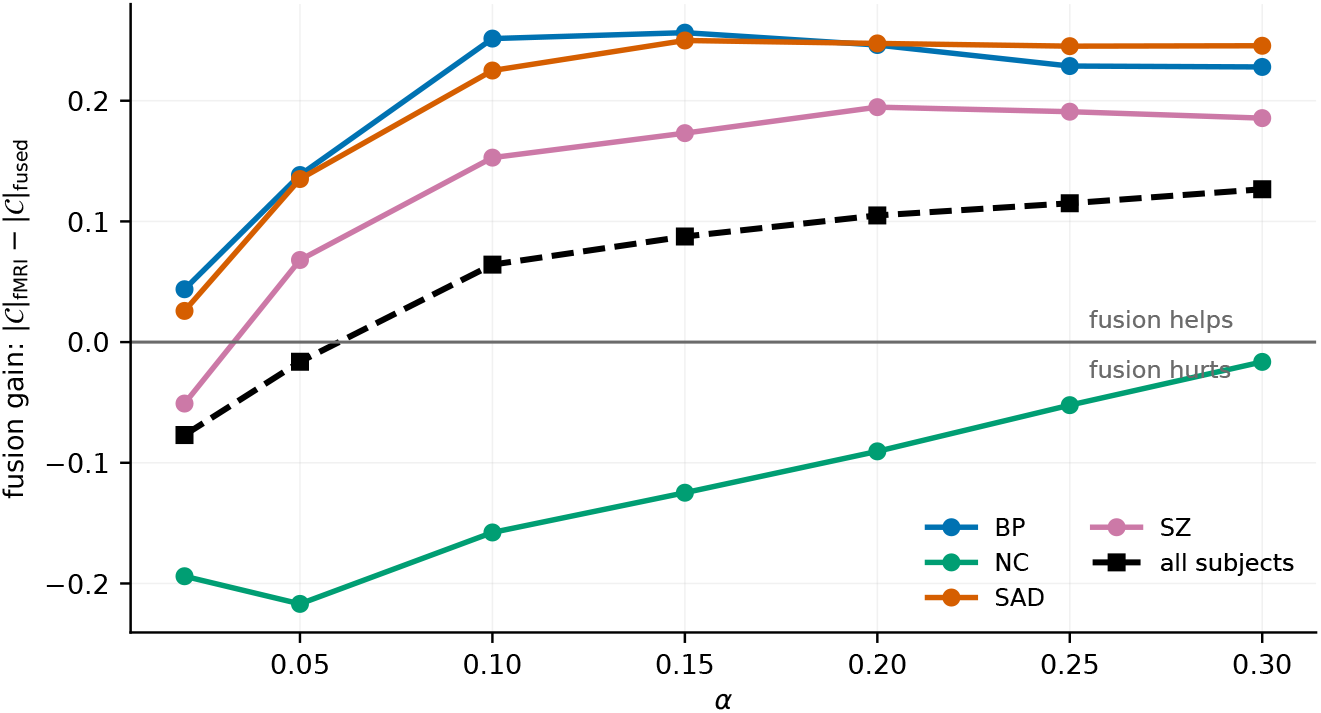
Fusion gain as a function of *α*. The sign is stable for BP, SAD and NC across the entire range; the aggregate (black, dashed) crosses zero between *α* = 0.05 and 0.10, so at strict confidence levels multimodal fusion is net harmful.

We note finally that the two unimodal prediction sets cannot simply be intersected: the intersection of two valid 90% sets attains only 82.1% coverage, and their union 98.3% (Jaccard overlap 0.62). Fusion at the model level, calibrated once, is the correct construction.

## 5 Discussion

On a four-way psychiatric classification task over 1,520 subjects, marginal split-conformal calibration attained 0.9000 empirical coverage against a nominal 0.90 while covering schizoaffective disorder at 0.819 and healthy controls at 0.941. The two deviations cancel, so the aggregate figure is uninformative about either. Class-conditional calibration reduced that 12.2-point spread to 0.4 points for an additional 0.07 labels of set size, under 3%.

The practical recommendation is narrow. Per-class coverage should be reported alongside the marginal figure, and calibration should be performed class-conditionally whenever diagnostic groups are unbalanced. On these data the second costs almost nothing in efficiency, so there is no efficiency argument for the marginal report. The failure mode we describe is not exotic: it requires only unbalanced classes and unequal separability, which together describe most clinical classification problems.

There is also a distributional point worth stating plainly. Under marginal calibration the excess coverage enjoyed by healthy controls is financed by a deficit on schizoaffective patients, so the group with the least diagnostic certainty receives the weakest guarantee, and the reported number gives no indication of it. Where conformal prediction is proposed as a safety mechanism for clinical deployment, this is the opposite of the intended behavior.

Beyond calibration, the prediction sets obtained under class-conditional coverage separate subjects into strata that admit a substantive interpretation. Because coverage is equalized across diagnoses, these strata cannot be explained by class prior or per-class accuracy, and their validation against an independent structural-MRI model indicates that they reflect properties of the subjects. The stratification is informative where a point prediction is not: it distinguishes subjects on which the model is confident from those lying on a specific diagnostic boundary, and both from a small group whose label the procedure does not recover at any operating point. The absence of confident schizoaffective cases, together with the concentration of bipolar and schizoaffective subjects in the ambiguous stratum, is consistent with the clinical difficulty of these categories and suggests that set-valued output is a more faithful summary of what neuroimaging supports than a forced single label.

Holding coverage fixed also makes set size comparable across representations. On this cohort the functional model leaves less residual ambiguity than the structural one, while their fusion helps only the intermediate diagnoses and harms controls, and the aggregate benefit of fusion changes sign with the confidence level demanded. This suggests that the question of whether to acquire a second modality is better posed per diagnosis, or per subject, than as a single aggregate comparison, and it points toward sequential acquisition policies in which a second modality is obtained only when the first leaves a subject ambiguous. Such policies require care, since naive optional stopping invalidates the coverage guarantee.

Several limitations bound these conclusions. Mean set size remains 2.65 of four labels at *α* = 0.10, so the sets quantify uncertainty rather than support a diagnostic decision, and we make no claim of clinical utility. Study and class composition are partially confounded in the pooled cohort, since FBIRN contributes no bipolar or schizoaffective subjects. The ambiguity marker used for external validation is model-derived and is not a clinical re-diagnosis; although it shares no predictions with the conformal procedure, it is not an independent clinical ground truth. The strata are defined from a modal set size across resamples, which is a per-subject summary rather than a single-split quantity. Finally, calibration and evaluation are performed on pooled sites; coverage under site shift, where exchangeability fails by construction, requires the weighted conformal treatment [8] and is left to future work. With 14 acquisition sites available, leave-one-site-out evaluation with covariate-shift weighting is the natural next step.

## Data Availability

FBIRN data are available through the Function Biomedical Informatics Research Network data repository [13]. B-SNIP data are available through the consortium under its data-sharing agreement [12].

## Ethics

FBIRN and B-SNIP data were collected under institutional review board approval at each contributing site, with written informed consent obtained from all participants.

## References

[1] V. Vovk, A. Gammerman, and G. Shafer, Algorithmic Learning in a Random World. Springer, 2005.

[2] J. Lei, M. G’Sell, A. Rinaldo, R. J. Tibshirani, and L. Wasserman, “Distribution-free predictive inference for regression,” J. Amer. Statist. Assoc., vol. 113, no. 523, pp. 1094–1111, 2018.

[3] M. Sadinle, J. Lei, and L. Wasserman, “Least ambiguous set-valued classifiers with bounded error levels,” J. Amer. Statist. Assoc., vol. 114, no. 525, pp. 223–234, 2019.

[4] Y. Romano, M. Sesia, and E. Candés, “Classification with valid and adaptive coverage,” in Adv. Neural Inf. Process. Syst., 2020.

[5] A. N. Angelopoulos, S. Bates, J. Malik, and M. I. Jordan, “Uncertainty sets for image classifiers using conformal prediction,” in Int. Conf. Learn. Represent., 2021.

[6] V. Vovk, “Conditional validity of inductive conformal predictors,” in Asian Conf. Mach. Learn., 2012.

[7] T. Ding, A. N. Angelopoulos, S. Bates, M. I. Jordan, and R. J. Tibshirani, “Class-conditional conformal prediction with many classes,” in Adv. Neural Inf. Process. Syst., 2023.

[8] R. J. Tibshirani, R. F. Barber, E. Candés, and A. Ramdas, “Conformal prediction under covariate shift,” in Adv. Neural Inf. Process. Syst., 2019.

[9] B.-S. Einbinder, S. Bates, A. N. Angelopoulos, A. Gendler, and Y. Romano, “Label noise robustness of conformal prediction,” J. Mach. Learn. Res., vol. 25, 2024.

[10] H. Rokham et al., “Addressing inaccurate nosology in mental health: a multilabel data cleansing approach for detecting label noise from structural magnetic resonance imaging data in mood and psychosis disorders,” Biol. Psychiatry Cogn. Neurosci. Neuroimaging, vol. 5, no. 8, 2020.

[11] Y. Du et al., “NeuroMark: an automated and adaptive ICA based pipeline to identify reproducible fMRI markers of brain disorders,” NeuroImage Clin., vol. 28, p. 102375, 2020.

[12] C. A. Tamminga et al., “Clinical phenotypes of psychosis in the Bipolar-Schizophrenia Network on Intermediate Phenotypes (B-SNIP),” Amer. J. Psychiatry, vol. 170, no. 11, pp. 1263–1274, 2013.

[13] D. B. Keator et al., “The Function Biomedical Informatics Research Network data repository,” NeuroImage, vol. 124, pp. 1074–1079, 2016.

